# Compressive stress drives morphogenetic apoptosis through lateral tension and Piezo1

**DOI:** 10.1101/2024.02.08.579454

**Authors:** Tatiana Merle, Martine Cazales, Thomas Mangeat, Ronan Bouzignac, Brice Ronsin, Christian Rouvière, Magali Suzanne

## Abstract

Tissues and organs are constantly submitted to physical stress, including compression, stretching, shear stress. The impact of compression due to overcrowding on cell extrusion has been the focus of recent studies. However, how tissue compression impact cell death in the context of morphogenesis is mostly unexplored. Here, we showed that a natural compression is exerted on the Drosophila developing leg by the surrounding tissue (or envelope) that is required for correct leg morphogenesis. In this tissue, apoptosis, preferentially localized in the future fold region, contributes to drive tissue folding through the generation of a pulling force on the apical surface. However, only a subset of these cells are dying within the expression domain of proapoptotic genes and how this precise pattern of cell death is established is totally unknown. We found that the natural compression exerted by the envelope contributes to the regulation of apoptosis, revealing that compression constitutes an integral part of apoptosis regulation during leg morphogenesis. We further reveal that compression drives a significant increase in lateral tension and favors apoptosis through the mechanosensor Piezo. Finally, perturbing cell cortex anchoring or membrane stiffness prove sufficient to block this process. Altogether, these results open new perspectives in term of mechanotransduction during morphogenesis.

## INTRO

Mechanobiology is a growing field of research addressing the contribution of physical parameters to living systems, including the physical properties of molecules, cells and tissues, the capability to generate and transmit forces, but also to respond to physical challenges such as shear stress, stretch or compression.

A number of pioneer studies in the field addressed the mechanical properties of isolated cells, revealing intriguing characteristic of cell mechanics, including how cell geometry can influence cortex tension and may control cell fate ^1^, the capability of cells to migrate in confined environment ^2,3^ or to adapt and respond to different environmental cues on bio-functionalized substrates ^4^. On the same line, *in vitro* studies on multicellular systems allowed to define how epithelial monolayers respond to different types of constraints including controlled stretching ^5^, or mechanical cues from their substrate ^6^.

It is now well recognized that tissue constraint is also an integral part of tissue homeostasis and morphogenesis. During morphogenesis, mechanics plays important functions as illustrated for example by the mechanical coupling between epidermis and muscle in c. elegans ^7^, the influence of tissue stretching on growth and oriented division ^8–10^, the activation of twist expression in the stomadeal primordium due to compression by the extension of the germ band ^11^. However, addressing the impact of physical constraint *in vivo* is still challenging due to some limitations regarding the possibility to probe and manipulate cells and tissues without perturbing their biological function.

Epithelial tissue, which constitute a protective barrier of organs from external cues, are particularly challenged and have to deal constantly with external constraint. In epithelial cells, mechanics also constitutes an integral part of fundamental cellular processes such as cell division, which has been extensively studied ^12,13^ or cell extrusion, which was shown to be sensitive to crowding *in vitro* ^14–16^. These works showed that, in monolayer epithelium, dead cell extrusion (or apoptotic cell extrusion) is triggered by singularities in cell alignments defined as topological defects in different epithelium types ^14^, and reveal that both apoptotic and live cell extrusion induced by overcrowding relies on Rho-kinase-dependent myosin contraction and the stretch-activated channel, Piezo ^15^. *In vivo*, cell extrusion was shown to be increased by crowding in the thorax of *Drosophila* ^17,18^. Thus, cell extrusion appears to be a way to buffer external constraints through the regulation of cell number and as such responds with high sensitivity to mechanical cues. More recently, a tight link between geometrical parameters (area and relative area) and the probability to enter apoptosis was established ^19^. This work shows that cells that just divided have a high probability to enter apoptosis and identified Hippo and Notch pathways as sensors of apical surface area or differential area between neighbors, in a fast-growing tissue. While these works revealed the importance of environmental cues or cell geometry to drive cell extrusion in highly proliferative tissues and overcrowding contexts, how these observations apply to morphogenesis is totally unknown.

Here, we use an original system to address the impact of compression on morphogenetic apoptosis, the developing leg of *Drosophila*, a tissue of cylindrical shape progressively elongating during metamorphosis ^20^. In this tissue, apoptosis is involved in the formation of stereotypic and concentric folds along the elongating leg through the generation of apico-basal forces. These forces favor the buckling of the epithelium at the level of a ring of cells with high level of apoptosis. We first show that this model is naturally under constraint through the compressive stress generated by the surrounding tissue, the peripodial epithelium or envelope. We further show that this compression contributes to fold formation and apoptosis regulation. Looking for factors that could favor apoptosis mechano-induction, we found that piezo plays a crucial role, while constraint induces dramatical changes in the subcellular distribution of membrane tension with a clear increase of lateral tension. Finally, perturbing moesin or alpha-spectrin function also blocks apoptosis mechano-induction highlighting the importance of cortex organization and/or membrane stiffness in constraint sensing. Based on these results, we propose that natural compressive stress drives an increase in lateral membrane tension in response to lateral membrane stretching, which results in the opening of piezo channel and apoptosis induction. These results point at an important role of lateral membrane in the perception of mechanical signals during morphogenesis.

## RESULTS

### I. Natural compression is required for leg morphogenesis

We first investigated whether the peripodial epithelium could exert compressive stress on the developing leg tissue as suggested by the frequent torsion of the leg during the elongation process. To test this hypothesis, we manually removed the peripodial epithelium at the beginning of metamorphosis or white pupal stage, before the formation of the most distal fold (t4-t5) (which we use as a model for the rest of this work) and compare the development of the leg in the presence or absence of this surrounding tissue (see schemes in Fig1A). Of note, this microdissection is coupled with a collagenase treatment, which help getting rid of the peripodial epithelium, without affecting the developing leg extracellular matrix (FigSup1). As soon as we removed the peripodial epithelium, we observed a sudden elongation of the tissue (see Fig1B, compare control (top) and without peripodial epithelium (bottom) conditions at t<20’), followed by a second, slower phase of elongation (see Fig1B and 1C for quantifications). This sudden release reveals the existence of compressive stress exerted by the peripodial epithelium on the developing leg epithelium, compressive stress that most probably increases during leg elongation due to the limit of extension of the peripodial epithelium.

**Figure 1:**
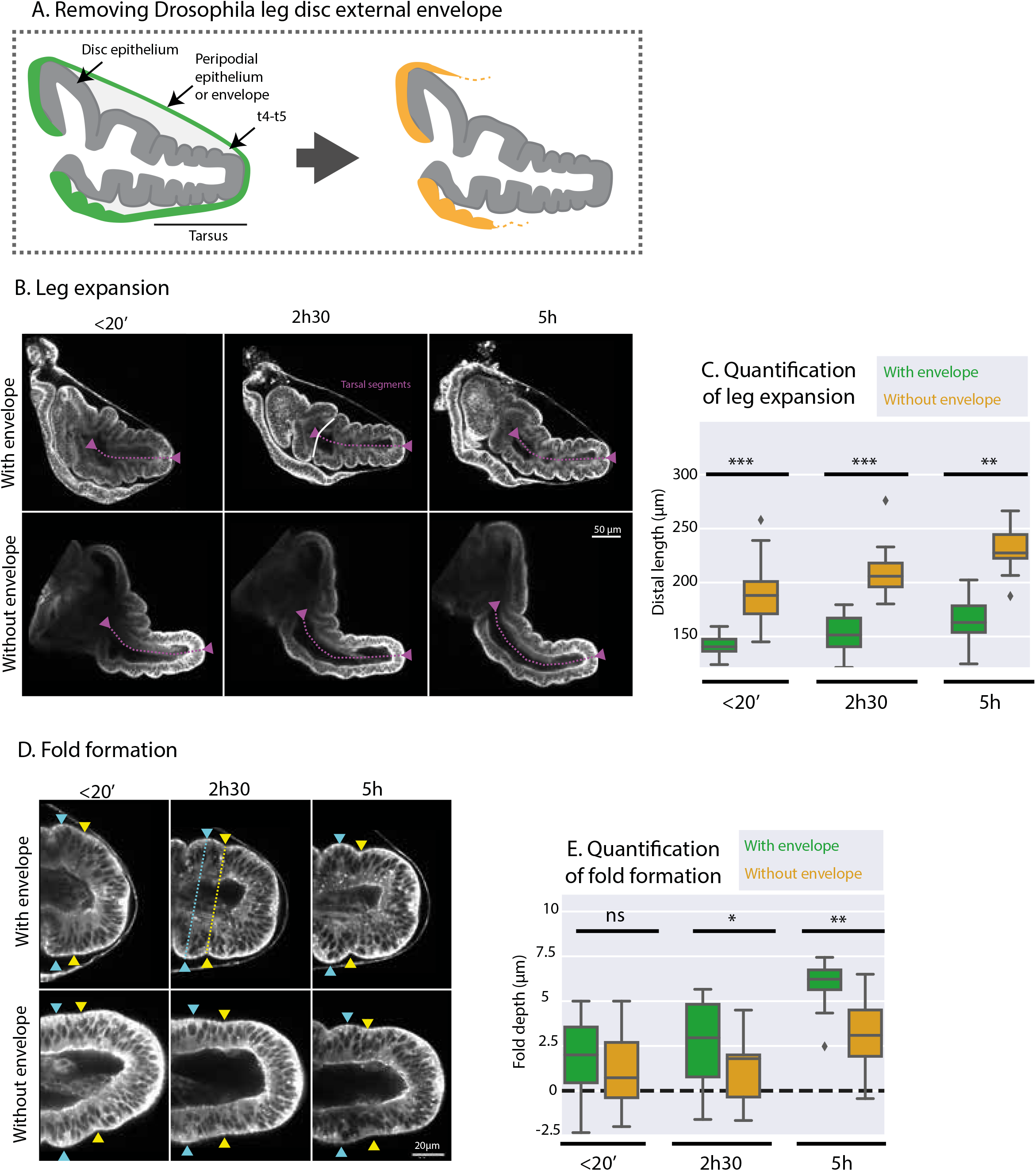
Compression is an integral component of morphogenesis. **A**. Scheme showing the general structure of the leg disc with the thick leg disc epithelium surrounded by the thin peripodial epithelium (PE) or “envelope”. PE is in green when unaltered, in yellow when microdissected. This colour code will be kept in all the figures. **B**. Still extracted from 2 movies of sqh-eGFP[KI] white pupae, showing the elongation of the leg disc in normal condition (top, with envelope) or without the envelope (bottom). **C**. Quantification of leg tarsus elongation with or without the envelope. The measured lengths from tibia-T1 to the tip of the leg is indicated are indicated in magenta in B. **D**. Fixed samples at different time point after dissection at white pupal stage and culture ex vivo, with or without the PE. The width of the segment is measured between the cyan triangles and the width of the fold is measured between the yellow triangles. **E**. Quantification of fold formation with (top) and without (bottom) PE as shown in D.

We then asked whether the release of this constraint could affect fold formation since all the folds formed in the elongating leg are formed before the opening and retraction of the peripodial epithelium ^20^. We observed that folds are less pronounced in the absence of external constraints, although some indentations can be seen after 5h of culture (Fig1D). However, these indentations never reached the depth of control folds in the absence of peripodial epithelium as shown by the fold depth quantifications (Fig1E, fold depth is determined by the difference between segment width and fold width). Overall, these observations indicate that drosophila leg develop naturally under natural compression and that this compression exerted by the peripodial epithelium on the elongating leg is required for proper morphogenesis of the tissue. Thus, external constraints constitute an integral part of leg morphogenesis.

### II. Compression influences the apoptotic pattern during morphogenesis

We next decided to decipher how constraints impact leg morphogenesis and fold formation. Since apoptosis was shown to constitute an important driving module of fold formation, generating an apico-basal pulling force that contributes to the increase of apical tension in the presumptive fold domain ^21^, we investigated how the apoptotic pattern evolves under different levels of constraint. To do so, we characterized the apoptotic pattern in the leg disc using a sensor of apoptosis (GC3Ai; Schott et al, 2017) expressed in the apterous domain (which includes the t4-t5 fold, see scheme in Fig2A) and allows the detection of apoptosis from its initiation to the fragmentation of the cell. We then tune external constraints either by removing the natural compression exerted on the elongating leg by dissecting the surrounding peripodial epithelium (no compression) or by increasing compression and preventing leg extension through the addition of agar in the culture medium (see schemes in Fig2B). We tried different concentrations and chose a concentration of agar preventing any further elongation of the developing leg. We then quantified the number of dying cells at different time points. Of note, we decided to exclude apoptotic bodies from our analysis, since these fragments (also positively marked by the apoptosensor GC3Ai) can subsist for several hours in the epithelium preventing the precise analysis of apoptotic dynamics, and quantified specifically early apoptotic cells (columnar cells positive for GC3Ai, see scheme in Fig2A). To test whether additional constraints can affect the number of apoptotic events, we cultivated the leg *ex vivo* for different periods of time and compared normal or additional compression conditions. While no significant difference was seen after 20’, the time required to dissect the leg discs and place them in culture, a clear difference can be seen after 1h. Of note, 1h is the time required for cells that started their apoptosis before the initiation of the experiment to be fragmented ^22^, thus the columnar dying cells counted after 1h correspond most probably to new apoptotic events. Interestingly, we observed an important increase of the number of dying cells with additional compression, compared to the control (Fig2C, D). This discrepancy is maintained at 2h30 and 5h. Altogether, these results suggest that constraint increases the number of apoptotic events. Reciprocally, we tested whether removing constraints affects the apoptotic pattern, by removing the peripodial epithelium and comparing the number of dying cells after 2h30 and 5 hours of culture. We chose this timing to avoid counting any false positive dying cells potentially due to the delay in the execution of apoptotic events started before dissection. We observed that when constraint is released, the number of dying cells drops by 75% after 2h30 (Fig2C, D). Then, after 5h, the number of apoptotic cells drops both in the control and in the “no compression” condition. These results suggest that compression is both necessary and sufficient to induce apoptosis in this developing tissue. It further supports the idea that compression is part of the regulation mechanism determining the pattern of apoptosis during this morphogenetic process.

**Figure 2:**
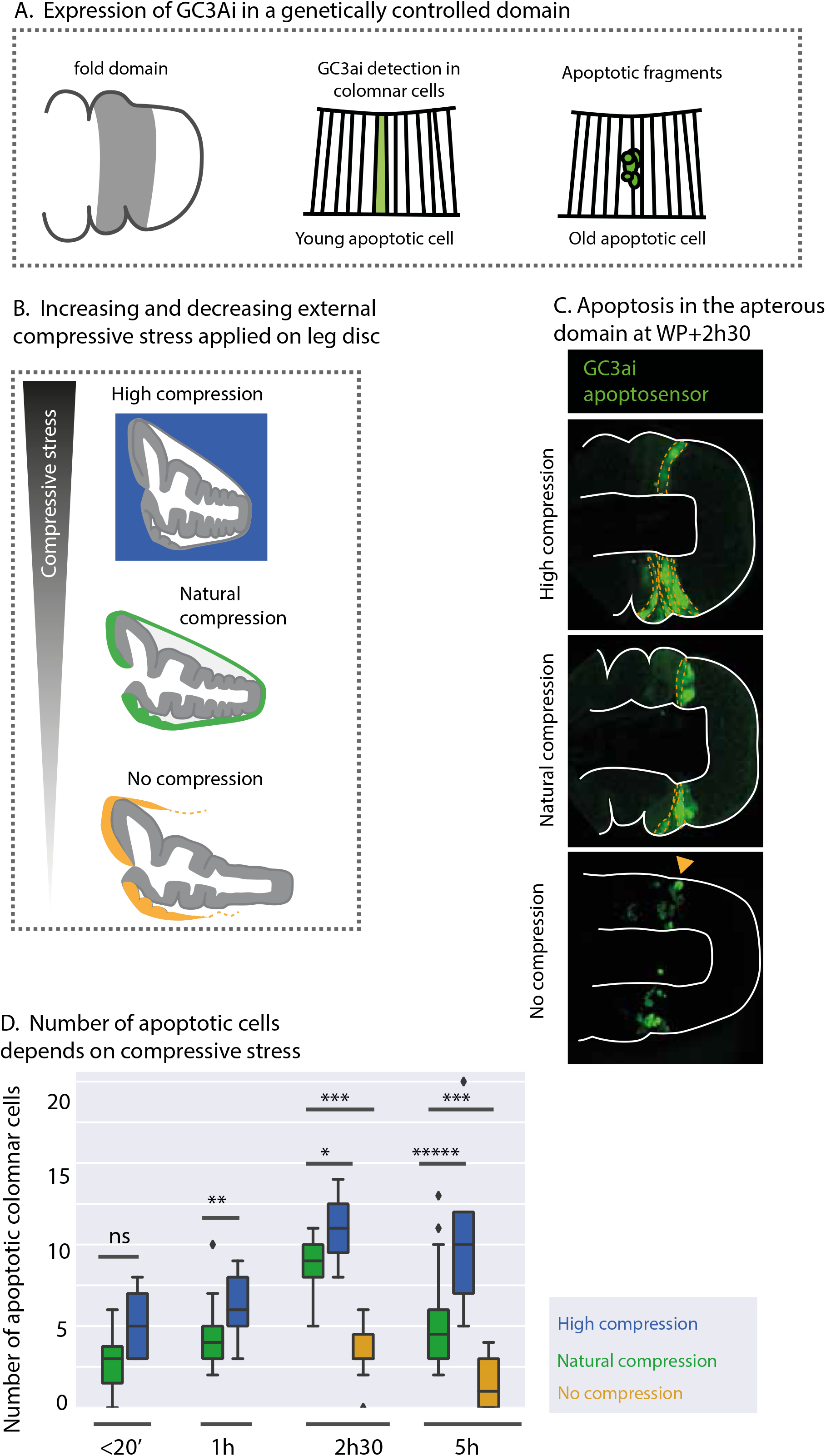
Natural compression drives morphogenetic cell death. **A**. (left) Scheme showing the expression domain of apterous (ap) in the leg disc, where the UAS-GC3Ai construct is expressed. (middle, right) Schemes showing that GC3Ai is detected both in early apoptotic cells (middle) and late apoptotic fragments (right). **B**. Schemes of the different conditions of tissue compression, going from high compression (top, blue color code), natural compression (middle, green color code) and no compression (bottom, yellow color code). **C. Cross sections** of apGal4; UAS-GC3Ai legs discs cultivated with different levels of compression (high, natural and no compression). Some apoptotic fragments are indicated by a yellow arrowhead, while early columnar apoptotic cells are outlined in yellow. **D**. Quantification of the number of early apoptotic cells observed in the different conditions shown in C.

### III. Impact of compression on apical cell surface and nuclear shape

Since apoptosis has been shown recently to be driven, at least in part, by geometrical parameters of cell apical surface ^19^, we decided to analyze how constraint affects apical cell geometry during leg morphogenesis. In particular, it was shown that apical surface area could help predict apoptosis activation in the thorax, while applying compressive stress on the developing tissue. Since leg tissue is a highly deformed tissue, we decided to use a recently developed segmentation tool adapted for automated segmentation in complex 2,5D structures called DISSECT (see Fig3A, Merle et al., 2023). Surprisingly, we found that cell apical apical area is mostly unchanged in the leg disc with or without natural compression (FigSup2A), strongly suggesting that this parameter is not key in apoptosis induction in this tissue.

**Figure 3:**
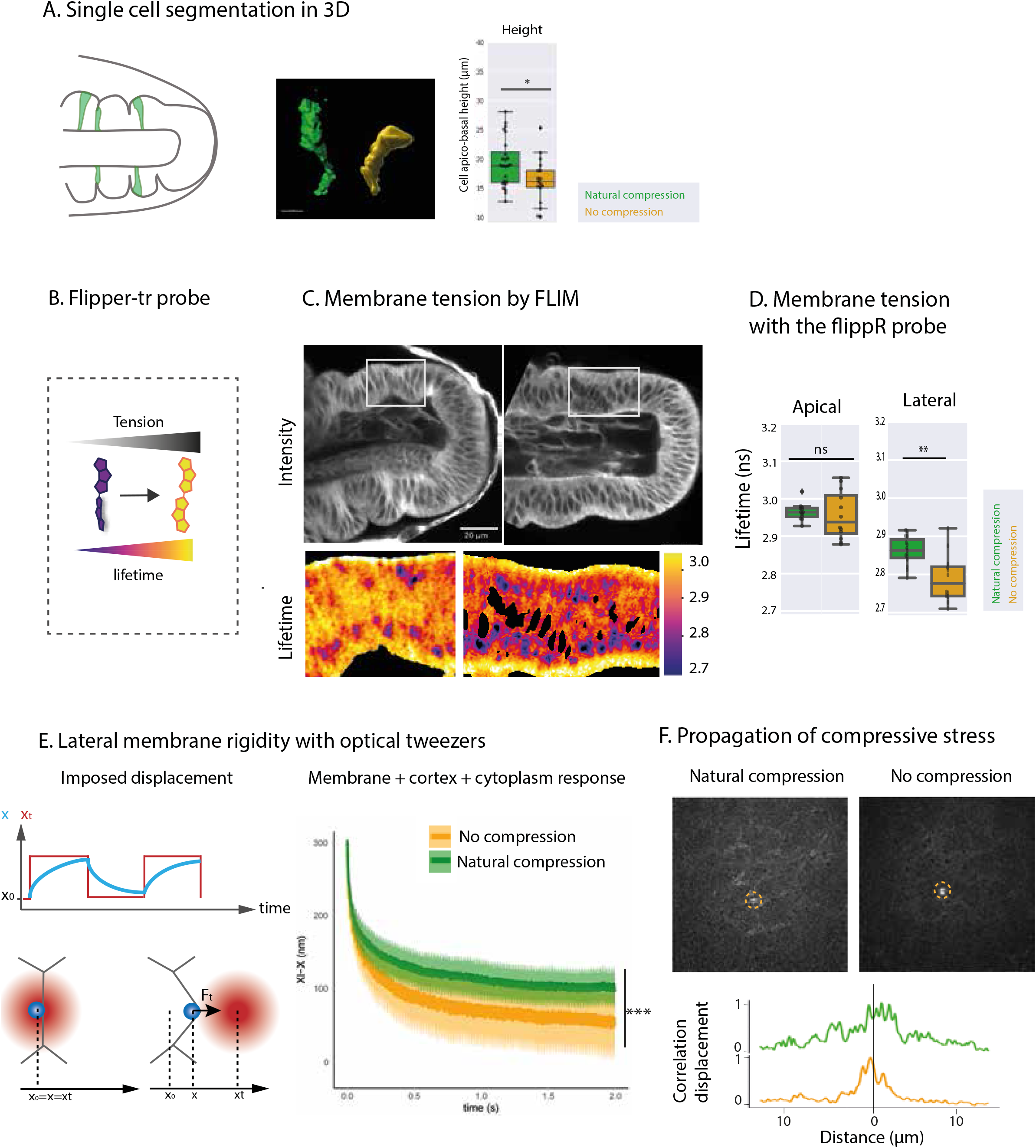
Impact of natural compression on cell geometry and membrane tension. **A. Quantification of cell geometry. A**. Scheme showing the distal part of the leg and showing the fold domain analyzed (left) and the readout obtained from z-stack of DE-cadh::GFP pupae 2.5h after puparium formation (APF) including automatized quantification of cell area using the Dissect program, with (green) or without (yellow) constraint from the PE. This color code is kept in all figures. **B. Cell segmentation in 3D. Left :** Scheme showing the principle of this single cell analysis through the random expression of UAS-lacZ construct in leg disc. **Middle** : Imaris 3D reconstruction of two cell in the fold region with and without compression at 2.5h after dissection. **Right**: Quantification single cell height in pupae with the following genotype: hsFlpG5, sqh-eGFPKI[29B]; UAS-LacZ; act-FRT-Stop-FRT-Gal4, UAS-HisRFP. **C**. Scheme from the Flipper probe showing it change of conformation under high membrane tension. **D**. Leg discs from white pupae cultivated for 2.5h with (left) or without (right) compression. Raw images (top) and life time color coded (bottom). **E**. Quantification of lateral membrane tension with (n=15) and without (n=11) compression. scale bar = 20um. **F**. Schemes showing the principle of optical twezers experiment. Lipid droplets are used as probes blue circle) and are pushed against the membrane to probe its resistance to deformation (bottom). The trap displacement is schematized in red, while the droplet displacement is shown in blue (top). The membrane is visualized with alpha-Spectrin::GFP, measures were done between 2 and 3h after dissection (WP ex-vivo culture). **G**. (Top) Optical tweezer experiment showing the displacement of membranes as a readout of force propagation in the tissue. The site of deformation is indicated by a yellow circle. Images before and after displacement are superimposed. Note that movement can be seen far from the manipulated region in the control (with compression, left), while it stays very local without compression (right). (Bottom) Curves of correlation displacement of membranes at different distances from the trap. Note that force propagate at a larger distance under constraint (green curve) compared to the local displacement with no compression (yellow curve).

Since no major changes were detected on the apical surface in response to the compressive stress applied to the developing leg tissue, we turned to a 3D characterization of the impact of compression. To investigate whether the nuclei were impacted by the natural compression exerted on the tissue, we segmented nuclei automatically using Cell Pose (FigSup2B) and extract nuclear aspect ratio. We observed that, the aspect ratio was significantly decreased when compression was removed (FigSup2C). These results indicate that the compression of the leg tissue by the peripodial epithelium is required to maintain the elongated shape of the nuclei observed in the control leg. This led us to suspect that the compressive stress could be sensed at the nucleus level, through this modification of nuclear shape.

The Hippo pathway is now well known to respond to mechanical forces ^13^. The translocation of Yorkie (readout of Hippo activation status) has also been shown to respond to changes in nuclear shape when cells are stretched in a rigid substrate ^24^. This pathway can also respond to crowding and is known to directly regulate cell survival and cell death through the activation of transcription of the apoptosis inhibitor by Yorkie ^25^. Since we observed a modification of nuclear shape under natural compression in the leg, we wondered whether the pattern of Yorkie nuclear translocation could be modified under different level of compression during the morphogenesis of the leg disc and be involved in apoptosis regulation in this tissue. To our surprise, the intensity of nuclear Yorkie was similar in the presence or in absence of compressive stress (with or without the compression exerted by the peripodial epithelium), strongly suggesting that the deformation of the nuclei observed under compressive stress is not sufficient to impact Yorkie import to the nucleus in this tissue (FigSup2D). To further test the implication of Yorkie in apoptosis induction in response to compressive stress, we expressed an RNAi against *yorkie* and analyzed the impact in term of apoptosis pattern with and without compression. To directly visualize the impact on compressive stress sensing, we divided the number of dying cells observed with natural compression by the number of dying cells observed in the absence of compression. This gives us the ratio of apoptotic cell number with and without constraint. We observed no significant changes in the apoptotic pattern when *yorkie* was inactivated in the leg tissue (FigSup2E). Indeed, we could not see any significative difference in the relative number of apoptotic dying cells (columnar cells expressing the apoptosensor). This is consistent with the fact that Yorkie distribution was unchanged in the leg disc with or without constraint.

Thus, the Hippo pathway do not seem to be involved in the regulation of apoptosis by compressive stress in this tissue and our results suggest that compression is controlling apoptosis by an independent mechanism.

### IV. Compression modifies the tension pattern and tissue dynamics

Given the radical change observed at the level of the nucleus in response to compressive stress, we reasoned that 3D cell shape could be also affected. Segmenting the whole cell in 3D, we found that cell height was significantly increase by the natural compression (Fig3A). This suggests that membrane tension could be redistributed in response to compression. We decided to characterize and compare membrane tension in the leg disc epithelium with or without natural compression. We used a live membrane tension reporter (a planarizable push-pull fluorescent probe called FliptR for fluorescent lipid tension reporter) that can monitor changes in membrane tension by changing its fluorescence lifetime ^26^ and use fluorescence lifetime microscopy (FLIM) to characterize membrane tension distribution (Fig3B, C). We found that apical tension was unaffected by compressive stress (Fig3D, left). However, lateral tension was strongly decreased when compression is removed (Fig3D, right). Indeed, lateral tension is clearly higher under compression compared to “no compression” condition, suggesting that the mechanical connection between apical and basal surface is favored by compressive stress. We further found that tissue contractility was modified under constraint. Indeed, we realized laser ablation in the depth of the epithelium and followed the repair process. While the tissue strongly contracts after damage in the control situation in order to seal the tissue (FigSup3A-B), when the tissue is free from external compressive stress, the contraction is far less important, revealing that overall lateral dynamics is strongly perturbed (FigSup3A-B).

Analyzing tension distribution with the FliptR probe, we noticed that while lateral tension was increase in the fold domain under compression, it is decreased in the distal domain of the claws (FigSup3D, compared to Fig3C). Interestingly, these two domains are submitted to different type of constraints by the peripodial epithelium: while the fold domain is submitted to a lateral compressive stress, the claw domain is submitted to an apical compressive stress, since compressive stress is applied at the tip of the leg (see scheme in FigSup4C). Since there is an important number of cells undergoing apoptosis in both domains, we asked whether this reciprocal modification of tension pattern under compression could lead to a reciprocal impact on apoptosis induction in these two domains. We thus compared the apoptotic pattern of the claw domain with and without peripodial membrane. We found that the apoptotic pattern was totally unperturbed in the claw domain by apical compression, as shown by the ratio of the number of apoptotic cells with and without compressive stress close to 1 (FigSup3E). We conclude that apoptosis in the claws is insensitive to compression, which suggests that the stress may not be sensed the same way in both domains.

We further analyzed the subsequent modification in term of cortex rigidity. We decided to probe the lateral membrane and test its resistance to deformation by optical tweezers (scheme in Fig3E, left panel). We trapped lipid droplets and used them as probes to deform lateral membrane. In this experiment, membrane resistance to deformation is translated into a bigger distance between the trap and the membrane (see scheme in Fig3F, left). Interestingly, we observed that the cortex was much more deformable in the absence of compressive stress, while it became more rigid upon compressive stress, as shown by the smaller distance between the trap and the membrane at equilibrium in the absence of compression (Fig3E, right panel). These results interestingly point at a strong modification of lateral tension and dynamics in the leg tissue by compressive stress.

We further analyzed how this increase in lateral tension impacts force transmission across the tissue. Here again, we used optical tweezers. This time, we induced local deformation at the level of the lateral membrane and analyze how far the deformation is transmitted on distanced membranes within a cross section of the tissue (Fig3F, top). Interestingly, we could observe that under compressive stress, local deformations are more efficiently transmitted along the tissue, while in the absence of compression, deformation stays local, supporting the fact that forces should propagate more efficiently in this condition. Indeed, in the control situation (natural compressive stress), a local deformation induces a deformation of membranes at several cell diameters from the trap (Fig3G, bottom).

### V. Morphogenetic apoptosis is driven by lateral compressive stress through piezo sensor

Our results point at a strong impact of compressive stress on lateral membrane tension, suggesting that compressive stress could be sensed by lateral membrane stretching. To test this hypothesis, we used two different ways to perturb cell cortex: perturbing moesin function, an anchor of the cytoskeleton to the plasma membrane by the expression of a dominant negative form (Moe-TD), or perturbing alpha-spectrin, a protein ensuring membrane rigidity through the formation of a network below the plasma membrane by the expression of an RNAi. Interestingly, in the Moe-TD context, while cells conserve a relatively normal shape under compressive stress, cell shape is totally perturbed in the absence of peripodial epithelium (Fig4A). This can also be observed in alpha-spectrin RNAi context, although at a much smaller scale. As a result, cells become less elongated or even cuboidal in the Moe-TD context when the compressive stress is absent (Fig4A). Thus, the columnar shape of the leg disc epithelium would be maintained by the cooperation between cell cortex rigidity and lateral compressive stress during fold formation. In addition, both genetic mutant contexts appear to inhibit constraint sensing. Indeed, they inhibit apoptosis mechano-induction, leading to a similar apoptotic pattern with and without constraint (Fig4B, D note that the ratio of apoptotic cell number with or without constraint is close to 1).

**Figure 4:**
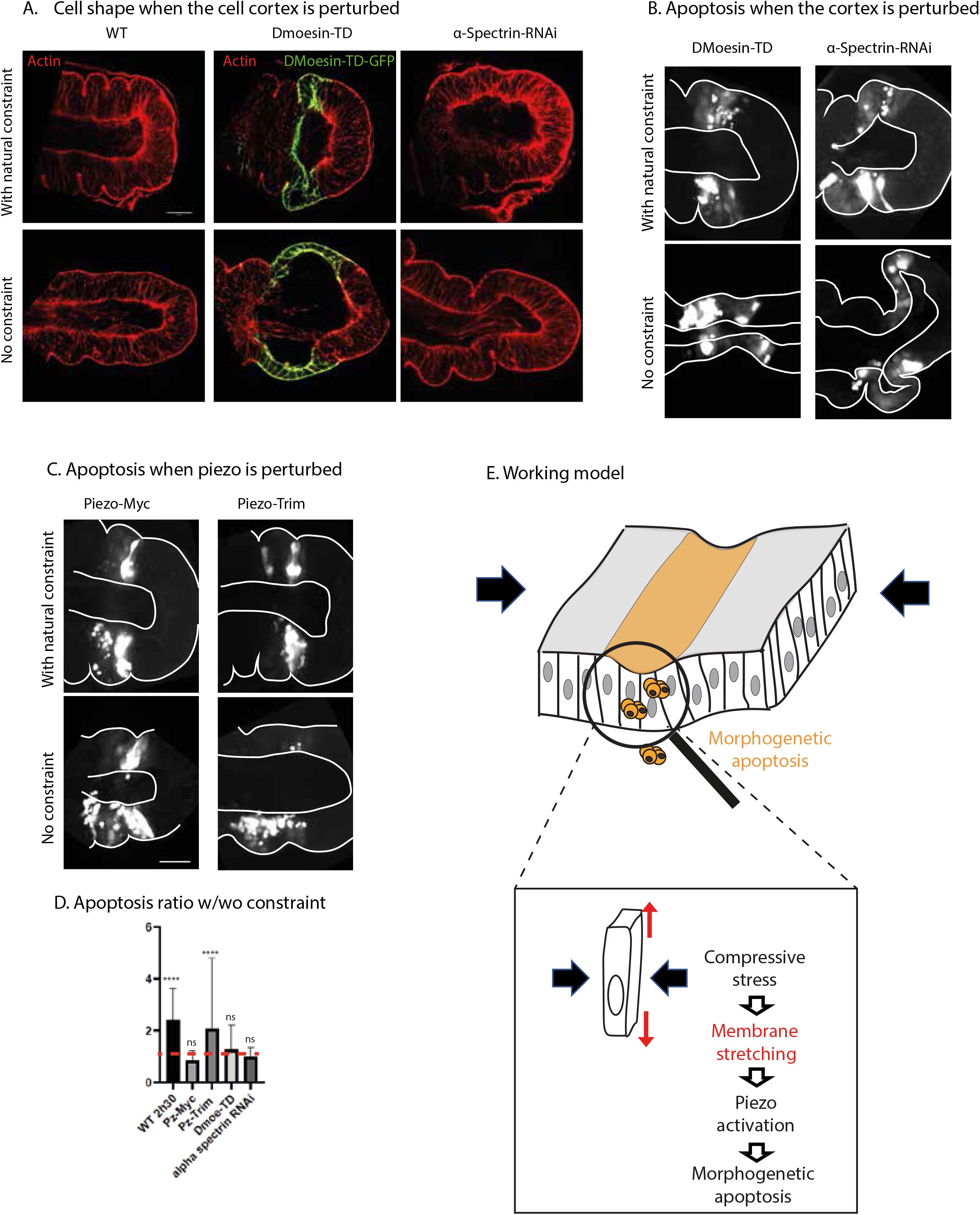
Molecular sensing of lateral compressive stress. **A**. Cross section of leg discs, 2.5h after dissection. F-actin is visualized in red in the whole panel. (Left) Control leg with (top) and without (bottom) compression. (Middle) apGal4; UAS DMoesin-TD-GFP leg discs with (top) and without (bottom) compression. (right) apGal4; UAS alphaSpectrin-RNAi leg discs with (top) and without (bottom) compression. scale bar = 20um **B-C**. Cross sections of apGal4;UAS-GC3ai leg discs, 2.5h after dissection with (top) and without (bottom) compression, expressing UAS-DmoesinTD (left) or UAS-alpha-spectrin RNAi (right) in B, or UAS-piezo-Myc (left) and piezo-TRIM (right) in C. **D**. Ratio of the number of apoptotic cells observed in the apterous domain with and without compression in the different genetical contexts presented in B and C. Piezo-Myc (n=17 and n=24), Pieozo-Trim (n=21 and n=17), DMoesin-TD (n=14 and n=12) and alphaSpectrin-RNAi (n=12)and n=13) with and without compression respectively. scale bar = 20um E. Working model showing the pro-apoptotic gene expression domain in yellow, in which apoptosis will be favored. Compressive stress induces lateral membrane stretching, piezo activation and finally drives morphogenetic apoptosis.

We next wonder what could be the sensor of this increase in lateral tension in the cell. Interestingly, it has been recently described that Piezo is distributed along the lateral cell membrane in epithelial columnar cells in the Drosophila wing disc ^27^. This led us to postulate that lateral membrane tension could be sensed by piezo and favor apoptosis as shown in overcrowding conditions ^15^.

To test whether inactivating piezo could affect the apoptotic pattern in the leg disc and modify the sensitivity to compressive stress, we expressed either a non-functional form of piezo (piezo-myc) or an activated form of piezo (piezo-TRIM) in the fold domain and analyzed the impact in term of apoptosis with and without the natural constraint. While the expression of piezo-TRIM has no visible impact on constraint sensing, we found interestingly that piezo-myc expression is sufficient to abolish the impact of compressive stress on the apoptotic pattern (Fig4C-D). Since piezo-myc encodes a non-functional channel, and since piezo homo-oligomerises to form the piezo channel, we suspect that this construct could have a dominant negative effect. This strongly suggests that a functional piezo channel is required to induce apoptosis in response to compressive stress.

## DISCUSSION

In this manuscript, we used as a model the Drosophila leg disc morphogenesis. We found that this tissue is under natural constraint during development. This constraint is a compressive stress exerted at the tip of the elongating leg by the peripodial epithelium. This compressive stress impacts lateral membrane tension. This increase in lateral tension is most probably the trigger of piezo channel activation and the subsequent morphogenetic apoptosis occurring in the fold domain and contributing to fold formation (Fig4E).

### Physical constraint as a driver of apoptosis during morphogenesis

A number of mechanosensitive pathways have been identified as driver of apoptosis in tissue homeostasis, where a balance has to be established between cell proliferation and cell death. Of note, a link between apoptosis and cell division has been made recently, showing that cell division can favor subsequent apoptosis ^19^. However, how this applies to a morphogenetic context where cell division is rare and cell change shape to form a new structure, is totally unexplored. Indeed, beyond the regulation of cell number, apoptosis plays also an important role in a number of morphogenetic process and can be actively involved in shaping tissue and organ ^28–30^. Here, we show that the drosophila leg disc is submitted to natural compression while a series of parallel folds are formed along the elongating tissue. This natural compression is exerted by the peripodial epithelium, a surrounding tissue that resist to the elongation of the leg. Indeed, we could observe an immediate release of the leg epithelium when the peripodial epithelium was removed by manual dissection. We further showed that this natural constraint participates to the regulation of apoptosis. While increasing compression is sufficient to increase the number of apoptotic events, removing compression strongly reduced the number of cells entering apoptosis. These results show that, during morphogenesis, the regulated pattern of apoptosis is tuned by natural compression. This imply that the natural compression exerted by the peripodial epithelium during leg extension is integrated in the developmental program of leg morphogenesis.

### The HIPPO pathway, the usual suspect for mechanically induced apoptosis

YAP (Yes-associated protein) and TAZ (transcriptional coactivator with PDZ-binding motif) are key mechanosensors and mechanotransducers controlling differentiation and proliferation in many cell types. YAP/TAZ nuclear localization can be affected by tissue mechanics in different ways, through cell-cell contact, cell–extracellular matrix contacts, cytoskeletal tension or cell shape ^13^. These different signals lead to the regulation of YAP/TAZ translocation to the nucleus. In fly tissues, the YAP/TAZ homologue Yorkie nuclear translocation is also known to respond to forces ^31–33^. The Hippo pathway is also well-known for its key role in the control of cell growth and apoptosis ^25^. Altogether, this make of Hippo a perfect candidate to sense compression in the leg disc and induce apoptosis in the forming fold.

However, here we found converging results strongly suggesting that he hippo pathway is unsensitive to the natural compression exerted on the developing leg. Indeed, while the leg disc is submitted to natural compression during its elongation process, as shown by the release of the tissue when the peripodial epithelium is removed, the level of nuclear yorkie remains mainly the same in the presence or in the absence of compression. In addition, the inactivation of yorkie has no impact on apoptotic pattern sensitivity to compressive stress, strongly supporting that a yorkie-independent mechanism is at play in this tissue.

### Apical constraint sensing

Recently, it was shown that compression leads to an increase in apoptosis through two complementary paths in the fly thorax: either a reduction of cell surface area or of relative surface area ^19^. These two paths were shown to depends on two signaling pathways: on the one hand, regulation of apoptosis through apical surface area depends on the Hippo pathway while on the other hand, the regulation of apoptosis through the relative area of a cell compared to its neighbors depends on the Notch pathway. We thus wonder whether apical surface area could be a parameter involved in constraint sensing in our system and driving apoptosis in the fold domain. However, apical surface appears unchanged between the compressed tissue and the non-compressed one, strongly suggesting that apical area is not a sensor of constraint in this tissue. If apical area and perimeter do not change much in response to compression in this tissue, we notice however that cell anisotropy was significatively different. Of note, a correlation has been made between anisotropy and high apoptotic rate previously in the thorax ^17^. However, this increase in cell surface anisotropy does not seem to induce any redistribution of apical tension, leading us to analyze 3D redistribution of tension in more detail.

### Tension redistribution under compressive stress

To analyze how compression impact global subcellular tension distribution, we used the FliptR tension probe. We did not observe any major changes in apical and basal tension comparing with and without peripodial membrane, however, we found that compressive stress has a strong impact on lateral membrane tension, which is significantly increased under the natural compressive stress exerted by the peripodial epithelium. Interestingly, piezo has been shown recently to be mostly localized at the level of lateral membrane in the columnar epithelium of the wing disc, this leads us to hypothesize that lateral membrane could be the main domain responding to lateral compressive stress.

To further test the role of lateral membrane in compressive stress sensing, we perturbed either cell cortex or cell membrane rigidity. Interestingly, these two approaches lead to similar results, both in term of cell shape and apoptotic pattern. In these two mutant contexts, columnar shape is only maintained is the presence of compressive stress. When the peripodial epithelium is removed, cells are no longer able to maintain their height and become less columnar or even cuboidal. Regarding the apoptotic pattern, it is no more sensitive to compressive stress in these mutant contexts, leading to a similar pattern in the presence or in the absence of compressive stress.

### Compressive stress could influence apoptosis dynamics

Apoptosis is a dynamic process relying on the activation of proteases named caspases. Caspase activation leads to the remodeling of cytoskeleton through the activation of ROCK and finally of Myosin II ^21,34–36^. In previous works, we have shown that apoptotic cells, while dying (before fragmentation), generate an apico-basal pulling force in the depth of the epithelium, through the formation of a linear structure of acto-myosin ^21,37^. Since we have shown here that natural compression influence Myosin stability and membrane tension pattern, it would be interesting to investigate in the next future, whether compression, beyond its impact on the induction of apoptosis, could impact apoptosis execution and dynamics. If Myosin II stabilization is favored by lateral compression, we could expect that apoptotic force would be more efficient to deform the apical surface of the epithelium and finally the tissue. It could also have an impact on apoptosis execution, since the apoptotic force is most probably also needed for the correct fragmentation of the cell and its proper elimination.

Finally, apoptotic cells also influence their neighbors, inducing, as a response to their own contractility, a ring of acto-myosin in the surrounding cells either to participate to their extrusion ^36,38–42^ or to contribute to morphogenesis ^28^. Since apoptosis is sensitive to compressive stress in this tissue, and since compressive stress favor force propagation, it is tempting to speculate that natural compression could also influence this non-autonomous impact of apoptotic cells and contribute to the propagation of apoptotic wave from ventral to dorsal in the fold domain.

Since apoptosis is a cellular process not only involved in morphogenesis, but also important in tumor development, acting either as an antitumoral or a pro-tumoral factor, it would be interesting to test how conserved is this pathway in a tumoral context.

## Acknowledgement

We thank Bruno Monier for constructive comments and suggestions along the project. MS’s lab is supported by grants from the National Agency of Research (ANR, PRC AAPG2021, CellPhy) and from the Research Association against Cancer (ARC, Programme Labélisé AAP2020, ARCPGA12020010001154_1591).

## Author contributions

**Tatiana Merle** conceived and performed the experiments in fly leg discs. She also participated in article writing. **Thomas Mangeat** realized and analyzed the optical tweezer experiments with the help **of Tatiana Merle. MC** set-up the conditions to cultivated leg disc with increased constraint and analyzed the apoptotic pattern in the different context of constraint. **RB** analyzed and quantified nuclei shapes using a home-made developed plug-in. **BR** help adjusting the set up for the acquisitions of the FLIM experiment. **CR** help develop Image J macro to segment nuclei. **MS** supervised the project, wrote the paper, and provided the funding.

## STAR Methods

### Experimental animals

The animal model used here is Drosophila melanogaster, in a context of in vivo/ex vivo experiments. In order to respect ethic principles, animals were anesthetized with CO2 (adults) before any manipulation. To avoid any release of flies outside the laboratory, dead flies were frozen before throwing them. Stocks of living flies were conserved in incubators, either at 18 or 25 degrees to maintain the flies in optimal condition. Fly food contains water, agar (0.8%), sugar (4%), flour (7.4%), yeast (2.8%), moldex (1%) and propionic acid (0.3%). Genotypes and developmental stages are indicated below. Experiments were performed in both males and females indifferently.

### Drosophila Stocks

E-cadherin-GFP (w; shg{TI}-eGFP; BDSC_60584) was obtained from Bloomington Drosophila Stock Center (BDSC). sqh-eGFPKI[29B], sqh-RFPtKI[3B] and UAS-GC3Ai were previously described (Ambrosini et al, 2019; Schott et al, 2017); as well as Yorkie_KI_GFP ^43^ and alpha-spectrin::GFP ^44^. Act-FRT-Stop-FRT-Gal4, His-RFP and ap^md544^-Gal4, were obtained from Bloomington. Clones were induced in individual with the following constructions: hsFlpG5, sqh-eGFPKI[29B]; UAS-LacZ; act-FRT-Stop-FRT-Gal4, UAS-HisRFP. DmoesinTD is from ^45^ ; piezo constructs from ^46^.

### Immunohistochemestry

Imaginal leg discs were dissected at white pupal stage in Schneider’s insect medium (Sigma-Aldrich) supplemented with 15 % fetal calf serum and 0.5 % penicillin-streptomycin as well as 20-hydroxyecdysone at 2 mg/mL (Sigma-Aldrich, H5142), incubated for 2,5 or 5h with or without 0,4% low melting agarose. Tissues were then fixed in 4% paraformaldéhyde (PFA 4%) for 20’, then extensively washed in PBT (PBS with 0.3% triton) and mounted in Vectashield (Vectors laboratories) or extensively washed in PBS-Triton 0.3%-BSA 1% (BBT) and incubated overnight at 4°C with appropriate dilutions of primary antibodies in BBT. Anti-E-cadherin antibody (Developmental Studies Hybridoma Bank – DCAD2) at 1:50 or anti-Lamin Dm0 (DSHB:ADL67.10 at 1:50). After washes in BBT, imaginal leg discs were incubated at room temperature for 2 h with 1:200 anti-rat IgG 647, anti-rat IgG 555, anti-rabbit IgG 555 or anti-mouse 555 at 1:100 (obtained from Interchim). Then, samples were washed in PBS-Triton 0.3%, suspended in Vectashield (Vectors laboratories) and mounted on slide.

### Ex vivo culture of leg imaginal disc

Imaginal leg discs were dissected at white pupal stage in Schneider’s insect medium (Sigma-Aldrich) supplemented with 10 % fetal calf serum and 0.6 % penicillin-streptomycin as well as 20-hydroxyecdysone at 2 mg/mL (Sigma-Aldrich, H5142). Leg discs were transferred on a slide in 12 mL of this medium (with or without 0,4% of low melting agarose) in a well formed by a 120 mm-deep double-sided adhesive spacer (Secure-SealTM from Sigma-Aldrich). A coverslip was then placed on top of the spacer. Halocarbon oil was added on the sides of the spacer to prevent dehydration.

### Peripodial epithelium removal

After dissection, leg discs were incubated with collagenase at a concentration of 1µg/mL, diluted in the culture medium and heated for 5min at 37°C. Gentle micro-dissections were performed during the incubation to break the external envelope and ensure total PE removal. The incubation in collagenase never exceeded 30min. We checked that this treatment did not affected the extracellular matrix of the leg epithelium (FigSup1).

### Confocal imaging

Samples were analyzed using a LSM880 confocal microscope fitted with a Fast Airyscan module (Carl Zeiss) and equipped with a Plan-Apochromat 40x/NA 1.3 Oil DIC UV-IR M27 objective. Z-stacks were acquired using either the laser scanning confocal mode or the High-Resolution mode (Airyscan). Airyscan Z-stacks were processed in ZEN software using the automatic strength (6 by default) and the 3D method.

### Laser ablation

Laser ablation experiments were performed using a pulsed DPSS laser (532 nm, pulse length 1.5 ns, repetition rate up to 1 kHz, 3.5 mJ/pulse) steered by a galvanometer-based laser scanning device (DPSS-532 and UGA-42, from Rapp OptoElectronic, Hamburg, Germany) and mounted on a LSM880 confocal microscope (Carl Zeiss) equipped with a 63x C-Apochromat NA 1.2 Water Corr objective (Carl Zeiss). Photo-ablation in the mid-plane of the apico-basal axis of the epithelium was done by illuminating at 90 % laser power during 5s along a line. Data analysis was performed with the ImageJ software to measure tissue height before and after repair.

### Quantification and statistical analysis

#### Nuclear segmentation

Nuclei segmentation was conducted based on immunofluorescence staining with an anti-Lamin Dm0 antibody using Cellpose with the cyto2 model, with a cell diameter set to 30 pixels and a flow threshold of 0.4. The masks were filtered using Label Size Filtering (Greater_Than; Size Limit (pixels): 8000) from the MorhoLibJ plugin in Fiji. The middle plane of each nuclei mask was extracted using a homemade macro in Fiji.

#### Nuclear aspect ratio

Nuclear aspect ratio and circularity were quantified using Label Analyzer (2D,3D) from the SCF-MPICBG plugin in Fiji. The assignment of nuclei to segment or fold regions was determined based on the X, Y position of the mass center of each nucleus.

#### Quantification of nuclear Yorkie

Utilizing the same image and segmentation parameters as those employed in the Nucleus geometry analysis, we employed Label Analyzer (2D,3D) from the SCF-MPICBG plugin in Fiji to quantify the mean nuclear signal of Yorkie (Yorkie_KI_GFP) from each nuclei mask in 3D. Then, the mean nuclear Yorkie signal was normalized by the mean grey level of the image.

#### Quantification of cell height

Clones were generated in individual of the following genotype: hsFlpG5, sqh-eGFPKI[29B]; UAS-LacZ; act-FRT-Stop-FRT-Gal4, UAS-HisRFP. Random expression of UAS-LacZ was generated by 15’ heat shock. The height of individual cell was measure using Imaris software.

#### Quantification of cell apical area

The open software DISSECT ^23^ was used to measure automatically this parameter.

#### Leg elongation

Legs lengths (T1 to distal tip) were measured with the ImageJ software.

#### Fold formation

We measured the difference between the width of the segment T4 and the width of the forming fold T4T5. To have the fold depth, the difference is divided by 2. Mann-Whitney tests were performed with the scipy python library.

#### Apoptosis pattern

Legs were observed with the Imaris software and only apoptotic columnar cells were counted (i.e. still attached apically and basally). Mann-Whitney tests were performed with the scipy python library.

#### Flipper probe

Flipper-TR probe was diluted in DMSO at 1mM concentration and stored at -20°C. Just prior the experiment, the probe was diluted to 1µM into the leg culture medium (Schneider + ecdysone). Legs were incubated 30min in the medium and not washed for FLIM acquisitions. We used a Leica SP8 Dive biphoton equipped with a FLIM module (Falcon) for experiments, we used a 970nm pulsed laser (DeepSee Spectra-Physics) at 80MHz. We collected the t1 lifetime. For analysis, with ImageJ, a filter was applied on the lifetime images: every pixel with an intensity (photon count, other image) smaller that 1000 was discarded. Mean measurements were done with ImageJ and Mann-Whitney tests were performed with the scipy python library.

### Optical Tweezers device

A home-built setup was used for optical trapping measurements. The system is a combination of a fluorescence microscope and an active optical trapping system. This trapping system has two components, one to precisely control the position of the optical trap on the sample and a back focal plane interferometric (BFPi) system to track the position of the trapped object relative to the center of the laser ^47,48^. For calibration and the measurements described below, a steering mirror conjugated to the pupil plane of the microscope (Thorlabs FSM75-P01) is used to achieve nanometric laser deflection. A 3 axes piezo electric stage (Piezo concept BIO3) is also used to precisely focus on the trap object. The optical trapping system is implemented in a conventional inverted fluorescence microscope (Leica DMI6000 B). The lens used is a 100x magnification lens with a numerical aperture of 1.4 (Leica HCX PL APO). To optimize the fill factor of the infrared laser at 85% of the objective lens pupil plane, an X4 afocal telescope consisting of two relay lenses is used (Thorlabs ACA254-050-1064 and ACA254-200-1064). The focused diffraction spot at the objective exit has a width of 600 nm at mid-height. This allows an efficient optical trap. The microscope is optimized for simultaneous GFP and RFP imaging combine with the infrared laser trap thanks to 3 band dichroic (Semrock Di03-R405/488/561/635-t1-25×36). Imaging and optical tweezers are achieved by combining a National Intruments acquisition card with the Inscoper synchronisation box. Inscoper software manages all the synchronisation steps.

### Optical Tweezers measurements and calibration procedure

The first step was to capture an internal lipid droplet (with an index close to 1.5 and an average size of 1 µm) and use it as a probe. It was then applied to a cell membrane at a depth of 5 µm from the apical surface of the cell. The laser used for trapping was centred at 1064 nm and the power injected into the sample was 200 mW. At this power and wavelength, the behaviour of the tissue is not affected. However, it is possible to displace the membrane cortex of the cells ^49^.

To visualise the force propagation in the junctions, a sinusoidal laser displacement was applied with a period of 0.12 Hz and an amplitude of 850 nm. At this speed, with respect to the damping coefficient of the medium, we remain within the linearity zone of the optical trap.

To measure the tension, a 320nm laser square step with a period of 0.12hZ was used for back focal plane interferometry tracking. The height of this step remains within the linearity of the position detector and allows us to calibrate the detector repose for each measurement in order to convert V into nm according to the size of the lipid droplet trapped. A Python program then averages 5 relaxation x curves for each measurement to reduce thermal or ATP fluctuations.

The fluctuation dissipation theory optical tweezers (FDT optical tweezers) force calibration method has been implemented to simultaneously calibrate the laser trap and the tracker conversion coefficient (V/nm) ^50,51^. In this method, when the properties of the optical medium are unknown or heterogeneous, the fluctuation dissipation theory at high frequency is used to calibrate the optical tweezers. A passive and active trajectory consisting of the highest frequency at 1HZ is recorded for 17 seconds to calibrate the optical trap for each measurement. The k-(trap constant spring), beta-(conversion coefficient V/nm) and elastic response of the medium are estimated simultaneously with an adapted program. The program is inspired by a multiplexed FDT method ^52^.

### Imaging processing for force propagation

First, commercial software (Scientific Volume Imaging, Huygens Pro) was used to deconvolve wide-field fluorescence images acquired during sinusoidal mechanical loading. Ten iterations with a signal-to-noise ratio of 4 were performed in the classical maximum likelihood mode.

Using the free software Fiji, the following protocol was followed. First, a bandpass filter (set from 12px to 3px) was used to extract the high frequency components of the image. The difference of the images per block of 10 images is cumulated 5 times to identify the cell membranes in the image field that move in a way that correlates with the movement of the optical trap. In fact, 10 frames is close to half a period of the applied sinusoidal stress. In this case, we are less sensitive to the fluctuations of the membranes that are naturally present throughout the image field. The images shown in Figure 3G are the projection of the maximum of previous displacement images of 28 measurements for the ctrl case and 33 measurements for the peripodial membrane case. A profile plot centered on the position of the optical trap in Figure3G is then normalized between 0 to 1 for each of the conditions. It is therefore possible to measure the long-range correlation of the mechanical strains induced by the optical trap.

A web application that calculates p-values by randomization was used to compare the plateau shown in Figure3F. The method is available here: http://huygens.science.uva.nl/PlotsOfDiff

## References

1. Fouchard, J., Bimbard, C., Bufi, N., Durand-Smet, P., Proag, A., Richert, A., Cardoso, O., and Asnacios, A. (2014). Three-dimensional cell body shape dictates the onset of traction force generation and growth of focal adhesions. Proc. Natl. Acad. Sci. U.S.A. 111, 13075–13080. 10.1073/pnas.1411785111.

2. Raab, M., Gentili, M., de Belly, H., Thiam, H.-R., Vargas, P., Jimenez, A.J., Lautenschlaeger, F., Voituriez, R., Lennon-Duménil, A.-M., Manel, N., et al. (2016). ESCRT III repairs nuclear envelope ruptures during cell migration to limit DNA damage and cell death. Science 352, 359–362. 10.1126/science.aad7611.

3. Thiam, H.-R., Vargas, P., Carpi, N., Crespo, C.L., Raab, M., Terriac, E., King, M.C., Jacobelli, J., Alberts, A.S., Stradal, T., et al. (2016). Perinuclear Arp2/3-driven actin polymerization enables nuclear deformation to facilitate cell migration through complex environments. Nat Commun 7, 10997. 10.1038/ncomms10997.

4. Ruprecht, V., Monzo, P., Ravasio, A., Yue, Z., Makhija, E., Strale, P.O., Gauthier, N., Shivashankar, G.V., Studer, V., Albiges-Rizo, C., et al. (2016). How cells respond to environmental cues – insights from bio-functionalized substrates. Journal of Cell Science, jcs.196162. 10.1242/jcs.196162.

5. Harris, A.R., Peter, L., Bellis, J., Baum, B., Kabla, A.J., and Charras, G.T. (2012). Characterizing the mechanics of cultured cell monolayers. Proc. Natl. Acad. Sci. U.S.A. 109, 16449–16454. 10.1073/pnas.1213301109.

6. Pérez-González, C., Ceada, G., Greco, F., Matejčić, M., Gómez-González, M., Castro, N., Menendez, A., Kale, S., Krndija, D., Clark, A.G., et al. (2021). Mechanical compartmentalization of the intestinal organoid enables crypt folding and collective cell migration. Nat Cell Biol 23, 745–757. 10.1038/s41556-021-00699-6.

7. Zhang, H., Landmann, F., Zahreddine, H., Rodriguez, D., Koch, M., and Labouesse, M. (2011). A tension-induced mechanotransduction pathway promotes epithelial morphogenesis. Nature 471, 99–103. 10.1038/nature09765.

8. Aigouy, B., Farhadifar, R., Staple, D.B., Sagner, A., Röper, J.C., Jülicher, F., and Eaton, S. (2010). Cell flow reorients the axis of planar polarity in the wing epithelium of Drosophila. Cell 142, 773–786. 10.1016/j.cell.2010.07.042.

9. Mao, Y., Tournier, A.L., Hoppe, A., Kester, L., Thompson, B.J., and Tapon, N. (2013). Differential proliferation rates generate patterns of mechanical tension that orient tissue growth. EMBO J 32, 2790–2803. 10.1038/emboj.2013.197.

10. Sagner, A., Merkel, M., Aigouy, B., Gaebel, J., Brankatschk, M., Jülicher, F., and Eaton, S. (2012). Establishment of global patterns of planar polarity during growth of the Drosophila wing epithelium. Curr Biol 22, 1296–1301. 10.1016/j.cub.2012.04.066.

11. Desprat, N., Supatto, W., Pouille, P.A., Beaurepaire, E., and Farge, E. (2008). Tissue deformation modulates twist expression to determine anterior midgut differentiation in Drosophila embryos. Dev Cell 15, 470–477. 10.1016/j.devcel.2008.07.009.

12. Irvine, K.D., and Shraiman, B.I. (2017). Mechanical control of growth: ideas, facts and challenges. Development 144, 4238–4248. 10.1242/dev.151902.

13. Panciera, T., Azzolin, L., Cordenonsi, M., and Piccolo, S. (2017). Mechanobiology of YAP and TAZ in physiology and disease. Nat Rev Mol Cell Biol 18, 758–770. 10.1038/nrm.2017.87.

14. Saw, T.B., Doostmohammadi, A., Nier, V., Kocgozlu, L., Thampi, S., Toyama, Y., Marcq, P., Lim, C.T., Yeomans, J.M., and Ladoux, B. (2017). Topological defects in epithelia govern cell death and extrusion. Nature 544, 212–216. 10.1038/nature21718.

15. Eisenhoffer, G.T., Loftus, P.D., Yoshigi, M., Otsuna, H., Chien, C.-B., Morcos, P.A., and Rosenblatt, J. (2012). Crowding induces live cell extrusion to maintain homeostatic cell numbers in epithelia. Nature 484, 546–549. 10.1038/nature10999.

16. Eisenhoffer, G.T., and Rosenblatt, J. (2013). Bringing balance by force: live cell extrusion controls epithelial cell numbers. Trends in Cell Biology 23, 185–192. 10.1016/j.tcb.2012.11.006.

17. Marinari, E., Mehonic, A., Curran, S., Gale, J., Duke, T., and Baum, B. (2012). Live-cell delamination counterbalances epithelial growth to limit tissue overcrowding. Nature 484, 542–545. 10.1038/nature10984.

18. Levayer, R., Dupont, C., and Moreno, E. (2016). Tissue Crowding Induces Caspase-Dependent Competition for Space. Current Biology 26, 670–677. 10.1016/j.cub.2015.12.072.

19. Cachoux, V.M.L., Balakireva, M., Gracia, M., Bosveld, F., López-Gay, J.M., Maugarny, A., Gaugué, I., di Pietro, F., Rigaud, S.U., Noiret, L., et al. (2023). Epithelial apoptotic pattern emerges from global and local regulation by cell apical area. Curr Biol 33, 4807-4826.e6. 10.1016/j.cub.2023.09.049.

20. Proag, A., Monier, B., and Suzanne, M. (2019). Physical and functional cell-matrix uncoupling in a developing tissue under tension. Development 146. 10.1242/dev.172577.

21. Monier, B., Gettings, M., Gay, G., Mangeat, T., Schott, S., Guarner, A., and Suzanne, M. (2015). Apico-basal forces exerted by apoptotic cells drive epithelium folding. Nature 518, 245–248. 10.1038/nature14152.

22. Schott, S., Ambrosini, A., Barbaste, A., Benassayag, C., Gracia, M., Proag, A., Rayer, M., Monier, B., and Suzanne, M. (2017). A fluorescent toolkit for spatiotemporal tracking of apoptotic cells in living. Development 144, 3840–3846. 10.1242/dev.149807.

23. Merle, T., Theis, S., Kamgoué, A., Martin, E., Sarron, F., Gay, G., Farge, E., and Suzanne, M. (2023). DISSECT is a tool to segment and explore cell and tissue mechanics in highly deformed 3D epithelia. Dev Cell, S1534-5807(23)00364-7. 10.1016/j.devcel.2023.07.017.

24. Elosegui-Artola, A., Andreu, I., Beedle, A.E.M., Lezamiz, A., Uroz, M., Kosmalska, A.J., Oria, R., Kechagia, J.Z., Rico-Lastres, P., Le Roux, A.L., et al. (2017). Force Triggers YAP Nuclear Entry by Regulating Transport across Nuclear Pores. Cell 171, 1397-1410.e14. 10.1016/j.cell.2017.10.008.

25. Huang, J., Wu, S., Barrera, J., Matthews, K., and Pan, D. (2005). The Hippo signaling pathway coordinately regulates cell proliferation and apoptosis by inactivating Yorkie, the Drosophila Homolog of YAP. Cell 122, 421–434. 10.1016/j.cell.2005.06.007.

26. Colom, A., Derivery, E., Soleimanpour, S., Tomba, C., Molin, M.D., Sakai, N., González-Gaitán, M., Matile, S., and Roux, A. (2018). A fluorescent membrane tension probe. Nature Chem 10, 1118–1125. 10.1038/s41557-018-0127-3.

27. Mim, M.S., Kumar, N., Levis, M., Unger, M.F., Miranda, G., Gazzo, D., Robinett, T., and Zartman, J.J. (2024). Piezo regulates epithelial topology and promotes precision in organ size control. Cell Rep 43, 114398. 10.1016/j.celrep.2024.114398.

28. Monier, B., and Suzanne, M. (2015). The Morphogenetic Role of Apoptosis. Curr Top Dev Biol 114, 335–362. 10.1016/bs.ctdb.2015.07.027.

29. Roellig, D., Theis, S., Proag, A., Allio, G., Bénazéraf, B., Gros, J., and Suzanne, M. (2022). Force-generating apoptotic cells orchestrate avian neural tube bending. Dev Cell 57, 707-718.e6. 10.1016/j.devcel.2022.02.020.

30. Toyama, Y., Peralta, X.G., Wells, A.R., Kiehart, D.P., and Edwards, G.S. (2008). Apoptotic force and tissue dynamics during Drosophila embryogenesis. Science 321, 1683–1686. 10.1126/science.1157052.

31. Ibar, C., Kirichenko, E., Keepers, B., Enners, E., Fleisch, K., and Irvine, K.D. (2018). Tension-dependent regulation of mammalian Hippo signaling through LIMD1. J Cell Sci 131, jcs214700. 10.1242/jcs.214700.

32. Kroeger, B., Manning, S.A., Fonseka, Y., Oorschot, V., Crawford, S.A., Ramm, G., and Harvey, K.F. (2024). Basal spot junctions of Drosophila epithelial tissues respond to morphogenetic forces and regulate Hippo signaling. Dev Cell 59, 262-279.e6. 10.1016/j.devcel.2023.11.024.

33. Rauskolb, C., Sun, S., Sun, G., Pan, Y., and Irvine, K.D. (2014). Cytoskeletal tension inhibits Hippo signaling through an Ajuba-Warts complex. Cell 158, 143–156. 10.1016/j.cell.2014.05.035.

34. Coleman, M.L., Sahai, E.A., Yeo, M., Bosch, M., Dewar, A., and Olson, M.F. (2001). Membrane blebbing during apoptosis results from caspase-mediated activation of ROCK I. Nat Cell Biol 3, 339–345. 10.1038/35070009.

35. Croft, D.R., Coleman, M.L., Li, S., Robertson, D., Sullivan, T., Stewart, C.L., and Olson, M.F. (2005). Actin-myosin-based contraction is responsible for apoptotic nuclear disintegration. J Cell Biol 168, 245–255. 10.1083/jcb.200409049.

36. Michael, M., Meiring, J.C., Acharya, B.R., Matthews, D.R., Verma, S., Han, S.P., Hill, M.M., Parton, R.G., Gomez, G.A., and Yap, A.S. (2016). Coronin 1B Reorganizes the Architecture of F-Actin Networks for Contractility at Steady-State and Apoptotic Adherens Junctions. Dev Cell 37, 58–71. 10.1016/j.devcel.2016.03.008.

37. Ambrosini, A., Rayer, M., Monier, B., and Suzanne, M. (2019). Mechanical Function of the Nucleus in Force Generation during Epithelial Morphogenesis. Dev Cell 50, 197-211.e5. 10.1016/j.devcel.2019.05.027.

38. Duszyc, K., Gomez, G.A., Lagendijk, A.K., Yau, M.-K., Nanavati, B.N., Gliddon, B.L., Hall, T.E., Verma, S., Hogan, B.M., Pitson, S.M., et al. (2021). Mechanotransduction activates RhoA in the neighbors of apoptotic epithelial cells to engage apical extrusion. Current Biology 31, 1326-1336.e5. 10.1016/j.cub.2021.01.003.

39. Kuipers, D., Mehonic, A., Kajita, M., Peter, L., Fujita, Y., Duke, T., Charras, G., and Gale, J.E. (2014). Epithelial repair is a two-stage process driven first by dying cells and then by their neighbours. J Cell Sci 127, 1229–1241. 10.1242/jcs.138289.

40. Ohsawa, S., Vaughen, J., and Igaki, T. (2018). Cell Extrusion: A Stress-Responsive Force for Good or Evil in Epithelial Homeostasis. Developmental Cell 44, 284–296. 10.1016/j.devcel.2018.01.009.

41. Rosenblatt, J., Raff, M.C., and Cramer, L.P. (2001). An epithelial cell destined for apoptosis signals its neighbors to extrude it by an actin- and myosin-dependent mechanism. Curr Biol 11, 1847–1857.

42. Teng, X., Qin, L., Le Borgne, R., and Toyama, Y. (2017). Remodeling of adhesion and modulation of mechanical tensile forces during apoptosis in Drosophila epithelium. Development 144, 95–105. 10.1242/dev.139865.

43. Fletcher, G.C., Diaz-de-la-Loza, M.-D.-C., Borreguero-Muñoz, N., Holder, M., Aguilar-Aragon, M., and Thompson, B.J. (2018). Mechanical strain regulates the Hippo pathway in Drosophila. Development 145, dev159467. 10.1242/dev.159467.

44. Khanna, M.R., Mattie, F.J., Browder, K.C., Radyk, M.D., Crilly, S.E., Bakerink, K.J., Harper, S.L., Speicher, D.W., and Thomas, G.H. (2015). Spectrin tetramer formation is not required for viable development in Drosophila. J Biol Chem 290, 706–715. 10.1074/jbc.M114.615427.

45. Polesello, C., Delon, I., Valenti, P., Ferrer, P., and Payre, F. (2002). Dmoesin controls actin-based cell shape and polarity during Drosophila melanogaster oogenesis. Nat Cell Biol 4, 782–789. 10.1038/ncb856.

46. Song, Y., Li, D., Farrelly, O., Miles, L., Li, F., Kim, S.E., Lo, T.Y., Wang, F., Li, T., Thompson-Peer, K.L., et al. (2019). The Mechanosensitive Ion Channel Piezo Inhibits Axon Regeneration. Neuron 102, 373-389.e6. 10.1016/j.neuron.2019.01.050.

47. Allersma, M.W., Gittes, F., deCastro, M.J., Stewart, R.J., and Schmidt, C.F. (1998). Two-dimensional tracking of ncd motility by back focal plane interferometry. Biophys J 74, 1074–1085. 10.1016/S0006-3495(98)74031-7.

48. Gittes, F., and Schmidt, C.F. (1998). Interference model for back-focal-plane displacement detection in optical tweezers. Opt Lett 23, 7–9. 10.1364/ol.23.000007.

49. Bambardekar, K., Clément, R., Blanc, O., Chardès, C., and Lenne, P.F. (2015). Direct laser manipulation reveals the mechanics of cell contacts in vivo. Proc Natl Acad Sci U S A 112, 1416–1421. 10.1073/pnas.1418732112.

50. Fischer, M., Richardson, A.C., Reihani, S.N.S., Oddershede, L.B., and Berg-Sørensen, K. (2010). Active-passive calibration of optical tweezers in viscoelastic media. Rev Sci Instrum 81, 015103. 10.1063/1.3280222.

51. Hendricks, A.G., Holzbaur, E.L.F., and Goldman, Y.E. (2012). Force measurements on cargoes in living cells reveal collective dynamics of microtubule motors. Proc Natl Acad Sci U S A 109, 18447–18452. 10.1073/pnas.1215462109.

52. Yan, H., Johnston, J.F., Cahn, S.B., King, M.C., and Mochrie, S.G.J. (2017). Multiplexed fluctuation-dissipation-theorem calibration of optical tweezers inside living cells. Rev Sci Instrum 88, 113112. 10.1063/1.5012782.

