## Supplementary material for "Compressive stress drives morphogenetic apoptosis through lateral tension and Piezo1": Fig S1-S3

Supplemental Figure 1

Figure Sup 1

The extracellular matrix of the leg epithelium is unaffected by the collagenase treatment

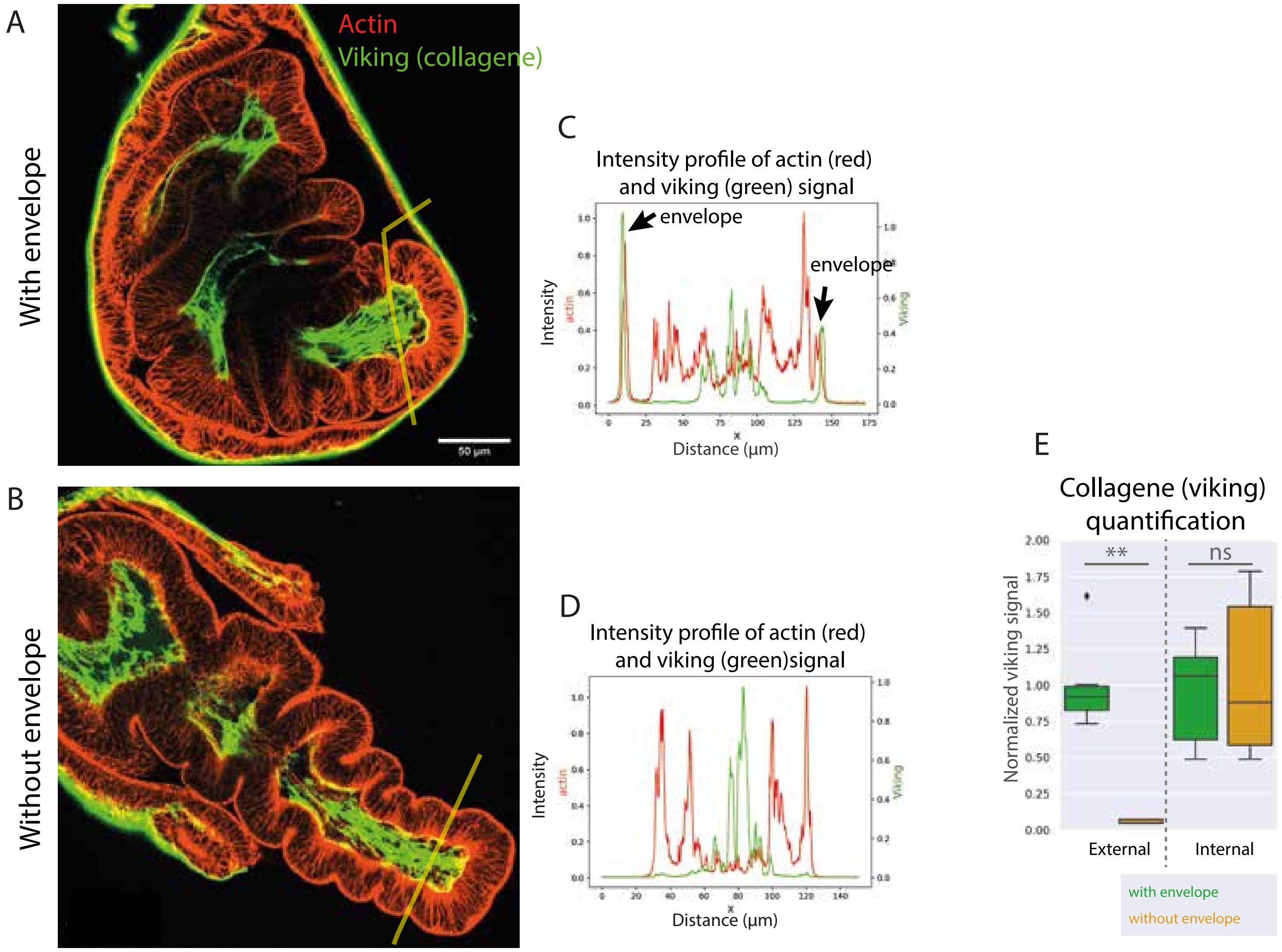

Supplemental Figure 2

Figure Sup 2

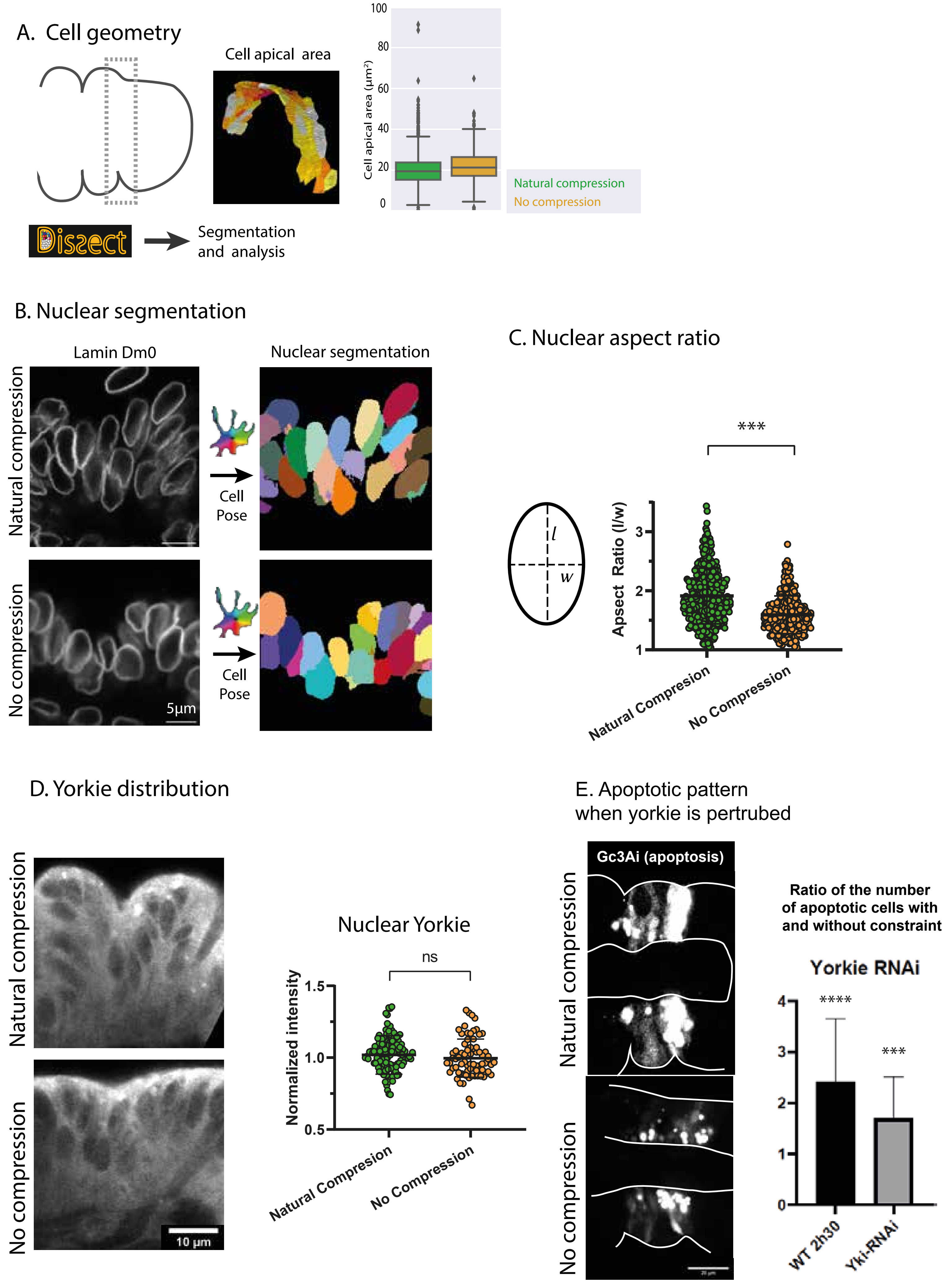

Figure Sup 3

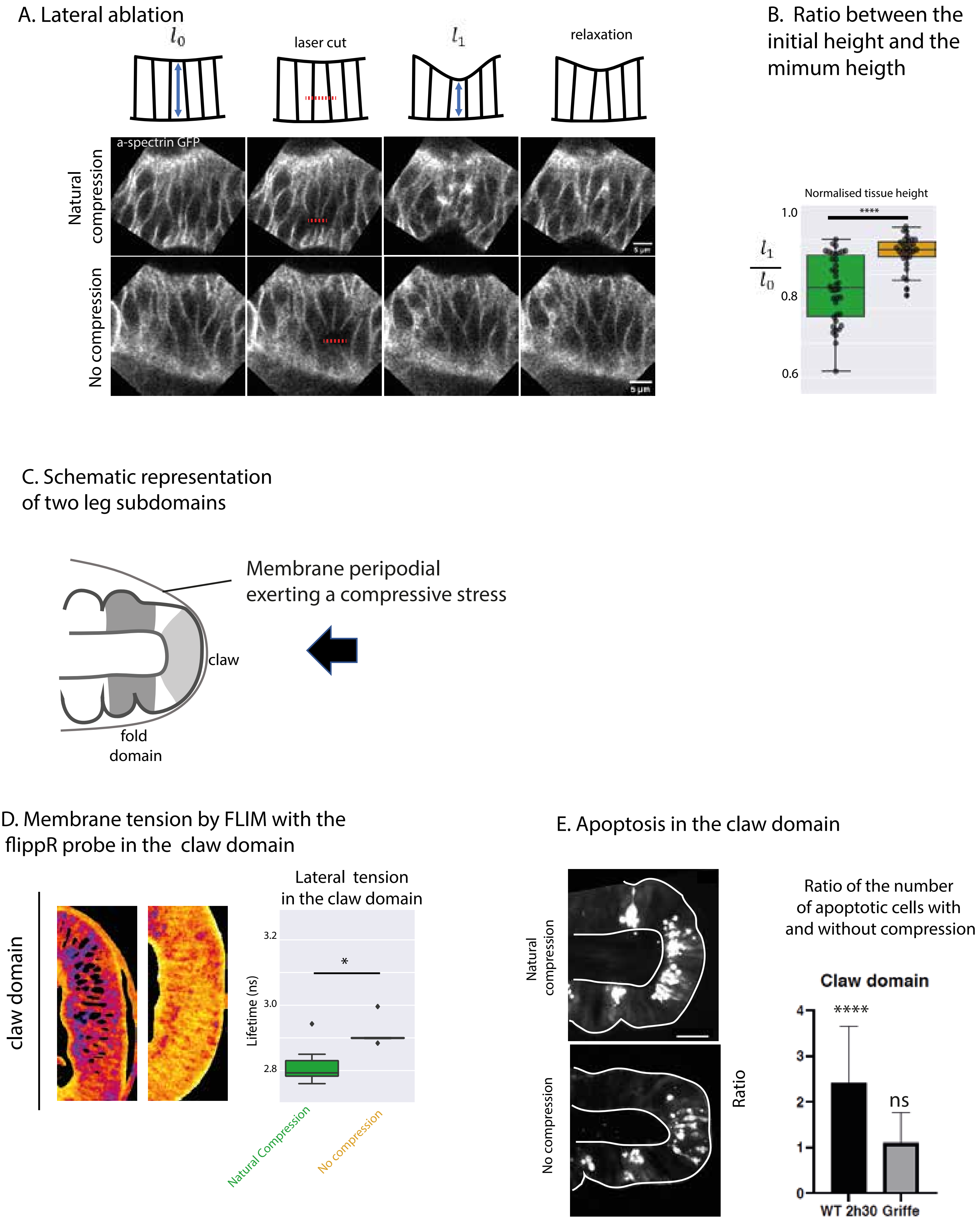
A. Lateral ablation
$l_0$
laser cut
$l_1$
relaxation
a-spectrin GFP
Natural compression
No compression
5 µm
B. Ratio between the initial height and the mimum heigth
Normalised tissue height
$\frac{l_1}{l_0}$
1.0
0.8
0.6
****
C. Schematic representation of two leg subdomains
Membrane peripodial exerting a compressive stress
claw
fold domain
D. Membrane tension by FLIM with the flippR probe in the claw domain
claw domain
Lateral tension in the claw domain
Lifetime (ns)
3.2
3.0
2.8
*
Natural Compression
No compression
E. Apoptosis in the claw domain
Natural compression
No compression
Ratio of the number of apoptotic cells with and without compression
Claw domain
Ratio
****
ns
4
3
2
1
0
WT 2h30
Griffe
